# An enhanced characterization of the human skin microbiome: a new biodiversity of microbial interactions

**DOI:** 10.1101/2020.01.21.914820

**Authors:** Akintunde Emiola, Wei Zhou, Julia Oh

**Affiliations:** The Jackson Laboratory for Genomic Medicine, Farmington, Connecticut, USA

## Abstract

The healthy human skin microbiome is shaped by skin site physiology, individual-specific factors, and is largely stable over time despite significant environmental perturbation. Studies identifying these characteristics used shotgun metagenomic sequencing for high resolution reconstruction of the bacteria, fungi, and viruses in the community. However, these conclusions were drawn from a relatively small proportion of the total sequence reads analyzable by mapping to known reference genomes. ‘Reference-free’ approaches, based on *de novo* assembly of reads into genome fragments, are also limited in their ability to capture low abundance species, small genomes, and to discriminate between more similar genomes. To account for the large fraction of non-human unmapped reads on the skin—referred to as microbial ‘dark matter’—we used a hybrid *de novo* and reference-based approach to annotate a metagenomic dataset of 698 healthy human skin samples. This approach reduced the overall proportion of uncharacterized reads from 42% to 17%. With our refined characterization, we revisited assumptions about the skin microbiome, and demonstrated higher biodiversity and lower stability, particularly in dry and moist skin sites. To investigate hypotheses underlying stability, we examined growth dynamics and interspecies interactions in these communities. Surprisingly, even though most skin sites were relatively stable, many dominant skin microbes, including *Cutibacterium acnes* and staphylococci, were actively growing in the skin, with poor or no relationship between growth rate and relative abundance, suggesting that host selection or interspecies competition may be important factors maintaining community homeostasis. To investigate other mechanisms facilitating adaptation to a specific skin site, we identified *Staphylococcus epidermidis* genes that are likely involved in stress response and provide mechanisms essential for growth in oily sites. Finally, horizontal gene transfer—another mechanism of competition by which strains may swap antagonistic or virulent coding regions—was relatively limited in healthy skin, but suggested exchange of different metabolic and environmental tolerance pathways. Altogether, our findings underscore the value of a combined reference-based and *de novo* approach to provide significant new insights into microbial composition, physiology, and interspecies interactions to maintain community homeostasis in the healthy human skin microbiome.

## BACKGROUND

Deep metagenomic shotgun sequencing is a powerful tool to interrogate composition and function of complex microbial communities. Microbial communities offer the potential for discovery of a tremendous suite of previously unknown biological functions, for example, new bioactive compounds, antimicrobials, virulence factors, or metabolic pathways. Such discovery has relied on the ability to survey and deconvolute species from mixed microbial consortia. Advances in next-generation sequencing and computational analyses have, in recent years, greatly furthered efforts to reconstruct microbial communities at the species^1,2^, strain^2,3^, and even single nucleotide polymorphism level^3,4^, examining function, transmission, and stability of the resident microbes.

However, interpretations of many metagenomic datasets are limited by the inability to characterize a large fraction of the total microbial reads present in the original sample^5,6^. This uncharacterized sequence space, or microbial ‘dark matter’^7^, typically results from the inability to map a sequence read to a known microbial reference genome and can exceed 96% of sequence reads within a sample^5^. Such ‘reference-based’ approaches, whether mapping reads to complete genomes^8^ or marker genes^9^, have high sensitivity and discriminatory ability between even very similar genomes^8^. However, microbes with no representative reference, or those with significant pangenomic variation, which can account for considerable within-species diversity in gene content^10^, are not captured. Conversely, reference-free approaches based on *de novo* assembly to aggregate reads into longer stretches of contiguous DNA sequence, can aid in the identification and characterization of new genomes. However, *de novo* assembly-based approaches are less effective in capturing small genomes (e.g., viruses), low-abundance microbes, and in discriminating between very similar genomes.

By combining both approaches into a holistic framework, we aimed to reduce the proportion of uncharacterized sequence space in a metagenomics dataset, and thus provide new insights into the biological function and interspecies interactions of these microbial communities. We used a hybrid *de novo* and reference-based approach aimed at characterizing microbial dark matter in the skin metagenome. Our previous analyses of this dataset (698 samples), which were exclusively reference-based, showed that the skin microbiome is defined primarily by the physiological characteristics of the skin site (e.g., whether it was a sebaceous, moist, dry, or foot site), then by host-intrinsic factors that confer individuality in strain representation and the presence of low-abundance and transient organisms^5,6^. More intriguing was the observation that the skin microbiome is remarkably stable even over years, despite the exposure of skin to different hygiene practices and the external environment^6^. However, our conclusions were based on an incomplete portrait with, on average, half of each sample remaining uncharacterized by our reference-based analyses^5^. By incorporating additional information from microbial dark matter, we stood to gain significant new insights into the landscape of skin biodiversity and microbial stability.

Leveraging our integrated approach, we uncovered previously unaccounted-for biodiversity and reduced microbial stability in the skin microbiome. We used this refined characterization to more deeply probe interspecies interactions, identifying intra-genus diversity and mechanisms underlying stability and inter-species interactions in the skin, including new assessments on growth rate and horizontal gene transfer. Our results demonstrate the highest resolution analysis of the skin microbiome to date, and provide new hypotheses for how skin microbes interact and compete to maintain homeostatic community conditions.

## RESULTS

### A hybrid *de novo* and reference-based microbial community analysis

To address the significant uncharacterized sequence space (mean ± sd 42% ± 24%) in our initial analysis of a 698-sample longitudinal skin metagenomic dataset (Supplementary Fig. 1), we used reference-independent approaches to reconstruct composition. With the improvement of *de novo* assembly algorithms to input large datasets^11^, we concatenated our samples and assembled iteratively, resulting in 75% ± 19% reads incorporated into the assemblies (Supplementary Fig. 2). The 1,037,465 resultant contigs >1kb were then grouped into genome ‘bins’ based on co-abundance clustering and nucleotide composition. However, because this approach is limited in its ability to recover small genomes, low-abundance species, and to ascertain precise taxonomic classifications, we speculated that integrating reference-based analyses (Fig. 1A) would further reduce dark matter beyond the 33% ± 21% reduction observed by mapping reads to our *de novo* reference set only (Fig. 1B). Microbial reads unmappable to our *de novo* reference catalogue were annotated by mapping to a reference database of fungal, bacterial, and viral genomes. Reference-based and *de novo* classifications were integrated with a normalization step that took into consideration the total proportion of reads derived from each approach (Supplementary Fig. 2). While using *de novo* references significantly aided reconstruction of microbiota, our hybrid approach most considerably reduced the proportion of dark matter (16% ± 17%; Fig. 1B and C).

**Figure 1.**
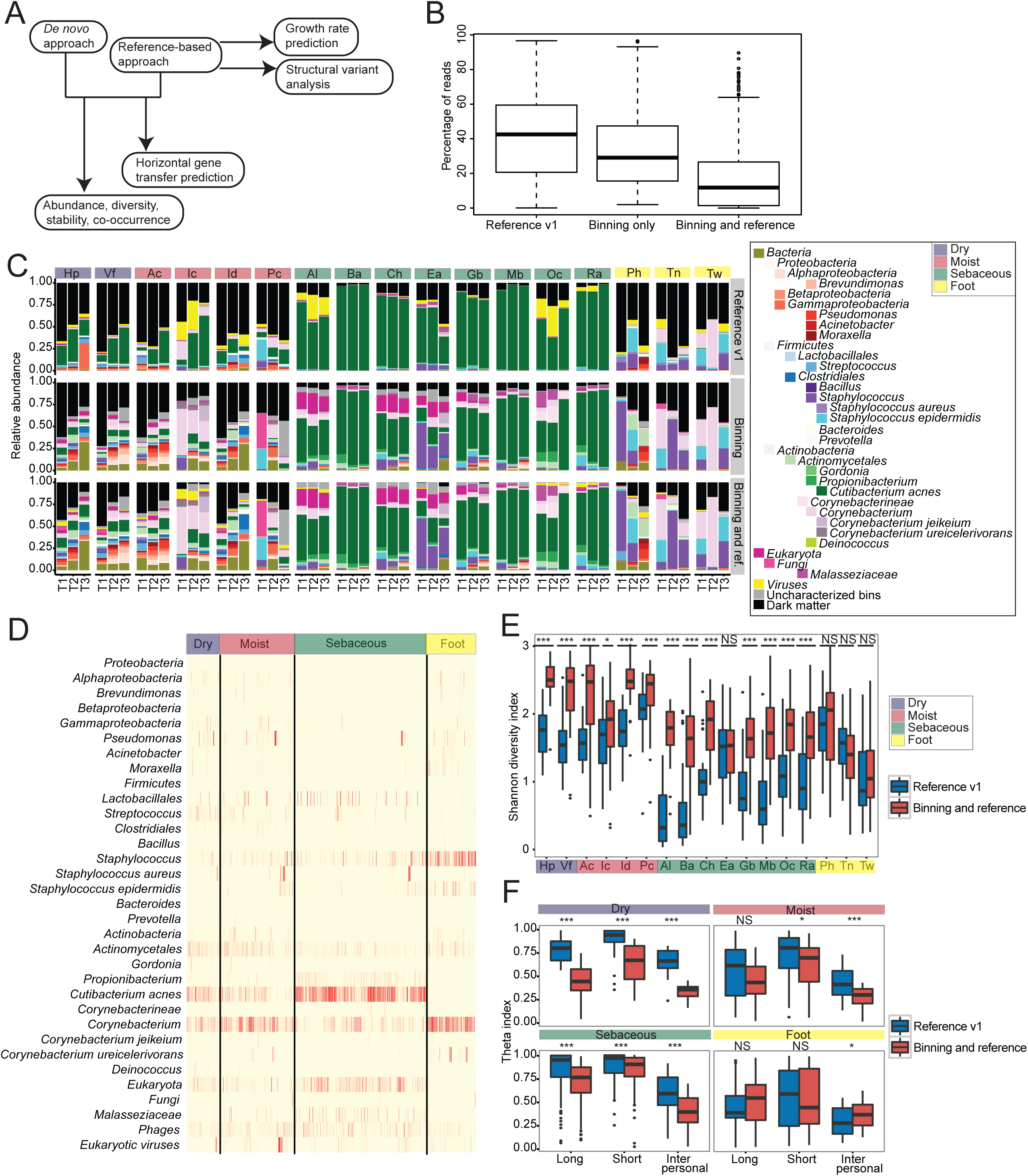
Hybrid *de novo* and reference-based approach resolves dark matter in skin metagenome. **(A)** Simplified flowchart of the hybrid *de novo* and reference-based characterization. **(B)** Boxplots show the fraction of uncharacterized sequences when using reference database from Oh *et al*.^6^, *de novo* genome bins only, or hybrid approach. Black lines indicate median; boxes first and third quartiles. **(C)** Microbial relative abundance across skin sites from a representative individual using the different classification approaches. **(D)** Heatmap shows microbial relative abundance across all samples from hybrid *de novo* and reference-based characterization, segregated by skin site characteristic. Darker colors indicate higher relative abundance. **(E)** Community diversity using Shannon diversity index. * = p-value < 0.05, ** = p-value < 0.01, *** = p-value < 0.001, NS = not significant by Wilcoxon-rank sum test. **(F)** Estimation of community stability using Yue-Clayton theta index, where θ∼1 represents a completely stable community. “Long” refers to sampling time interval between T1 and T2 while “Short” represents short sampling time interval between T2 to T3. * = p-value < 0.05, ** = p-value < 0.01, *** = p-value < 0.001, NS = not significant by Wilcoxon-rank sum test.

### A new biodiversity of the human skin metagenome

Our new compositional analysis was largely concordant with our previous findings that the skin microbiome is predominated by *Staphylococcus, Cutibacterium* (formerly *Propionibacterium*), *Corynebacterium*, and *Malassezia* species^5,6^ (Fig. 1C, Fig. 1D). However, we uncovered considerably more diversity of *Staphylococcus, Corynebacterium, Proteobacteria*, and fungal genomes at most skin sites (Fig. 1C, Supplementary Fig. 3A). For example, we identified a previously uncharacterized, but abundant, *Lactobacillales* colonizing the external auditory canal (Ea) as *Alloiococcus otitis*. 9% ± 13% of reads mapped to bins were unclassifiable based on BLASTn alignment to the NCBI nt database. These likely represented contigs that belong either to uncharacterized genomes or undiscovered pan-genomic variation of a lower abundance strain that could not be binned with its species unit. These otherwise unclassifiable bins were most frequently bacterial, underscoring that the majority of undiscovered biodiversity in the skin is not fungal or viral (Supplementary Fig. 3B).

Our revised classification showed lower representation of *C. acnes* than previous analyses, and markedly lower *Propionibacterium* phage representation in sebaceous regions. *De novo-*only approach did not capture viral contigs; these were most accurately recovered with the combined use of reference genomes, for example in the alar crease (Al) (Fig. 1C).

With our integrated classification, we re-examined our conclusions of diversity and stability at the community level. All skin sites except the ear and foot had higher diversity than previously reported, which is expected given the identification and inclusion of more genomes/genome bins than in our original analyses (Fig. 1E). Because resolved dark matter only represented a few genomes in the external auditory canal (Ea) (mostly *Lactobacillales*) and foot sites (mostly *Staphylococcus* and *Corynebacterium*), diversity was unchanged in these regions. Since previous analyses correlated increased diversity with decreased stability^6^, we re-evaluated our conclusions on stability over the ∼month (T2-T3) and ∼year intervals (T1-T2) collected in this study. Most sites were less stable than originally defined (Fig. 1F)—for example, the hand and forearm (dry) sites, which are highly exposed to the external environment (Fig. 1F, Supplementary Fig. 4A). This may be due to behavioral patterns like hand-washing or increased acquisition of transient, environmental microbes. Longitudinal tracking of individual species’ dynamics over time showed that community stability is driven by specific microbes (Supplementary Fig. 4B). For example, newly identified *Corynebacterium* species (e.g., *jeikeium*) were associated with stability whereas staphylococci showed more fluctuation over time.

### Mechanisms underlying interspecies interactions: growth dynamics of skin microbiota

A significant limitation of previous skin microbiome studies is the absence of information on microbial activity, which is necessary to understand potential underlying homeostatic mechanisms as microbes are unlikely to be completely inactive in the skin. We reasoned that the contributions of viable populations to overall microbial abundance and functional community potential could be inferred by assessing bacterial growth rate from the skin metagenome. This would also allow us to estimate the ratio of rapidly growing to dead/stationary cells of common skin microbes, which would provide additional insights into antagonistic interspecies interactions.

To achieve this, we used the peak-to-trough ratio (PTR) method^12^, implemented in GRiD^13^, which maps metagenomic reads to a bacterial reference genome to calculate coverage drops across the genome. Because most bacteria harbor a single circular chromosome replicated bi-directionally from the origin of replication (*ori*) to the terminus (*ter*) region^14^, rapidly growing cells will have a higher coverage at *ori* vs *ter*.

First, we generalized our analysis to define the steady-state growth dynamics of dominant skin microbes. Defining a microbe with a GRiD score > 1 as being in exponential phase, we identified the proportion of bacteria that were actively growing across different skin sites. Most examined microbes in dry sites (i.e., palm and forearm) were most active (Fig. 2A), interesting because these regions are often perturbed (e.g., by hand washing) and biomass is low. Such factors could affect microbial viability and growth rate to replenish the endogenous community. In contrast, sebaceous sites, which typically harbors higher biomass, least supported rapid growth, which could reflect the relatively specialized physiologic growth conditions, including anoxia, which would limit or slow growth of many microbes^15^. Foot sites favored growth of *S. epidermidis* and other non-lipophilic microbes at the expense of sebum-metabolizing *C. acnes*, which was least active at that site (Fig. 2A). Strikingly, growth rates of these microbes were stable over multiple timepoints (Fig. 2B).

**Figure 2.**
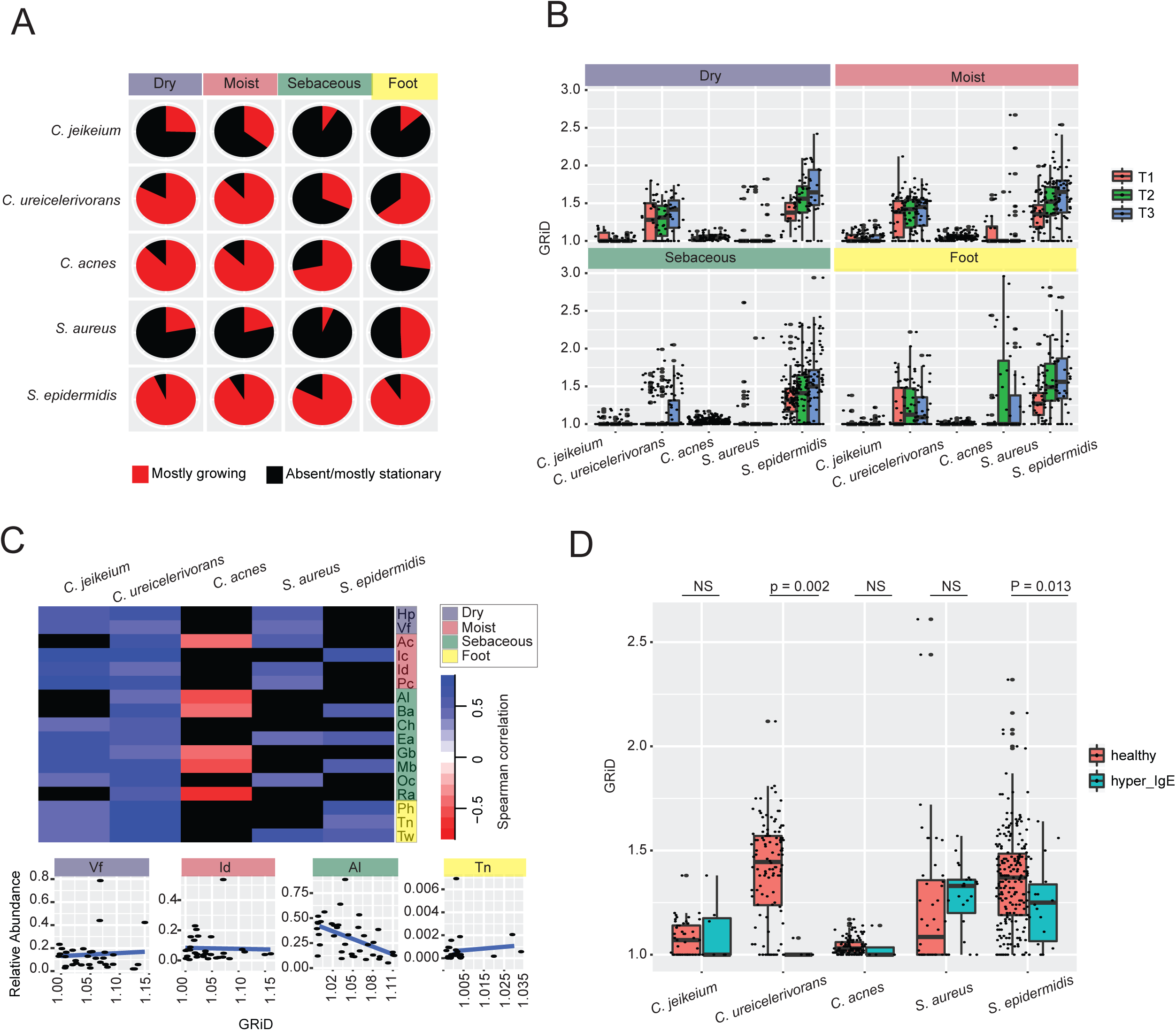
Growth dynamics of skin bacteria. Growth rate was calculated using GRiD^13^. **(A)** Proportion of common skin microbes in exponential vs. stationary growth phase/absent across different skin sites. GRiD = 1 for a microbe was considered stationary phase. *C. acnes= Cutibacterium acnes, C*=*Corynebacterium, S*=*Staphylococcus.* **(B)** Boxplot showing the GRiD score of microbes over time and grouped by skin site characteristics. **(C)** The heatmap shows partial Spearman correlation coefficient values, correcting for multiple measurements, between growth rate (GRiD) and relative abundance. Black colors indicate no significant correlation. The scatter plots below shows correlation between *C. acnes* growth rate (GRiD) and relative abundance from representative skin sites. **(D)** Boxplot showing microbial growth rate (GRiD) in healthy and primary immunodeficiency cohorts. Significant differences between groups were computed with Wilcoxon rank-sum test.

We also asked if increased growth rate resulted in increased relative abundance. Relative abundance was positively correlated with growth rate only for certain species (Fig. 2C). For example, *C. ureicelerivorans’* relative abundance was strongly correlated with growth rate in all skin sites (Fig. 2C). This suggests that for select species, microbial abundance may be controlled by how rapidly cells are dividing. In contrast, *C. acnes* relative abundance and growth rate were strongly anti-correlated in sebaceous sites (Fig. 2C), suggesting that *C. acnes*’ growth rate may be modulated based on the presence/absence of competition or is regulated by its quorum sensing mechanisms.

Finally, we examined the microbial growth dynamics in patients with primary immunodeficiency syndrome, characterized with eczematous lesions^5^, as rapid growth might identify potential pathogens. Common skin commensals such as *S. epidermidis* and *C. ureicelerivorans* had significantly decreased growth rate in these patients (Fig. 2D). The decreased growth rate of *S. epidermidis* is surprising because these patients are unusually prone to staphylococcal infections. This observation may be due to the lack of correlation between growth rate and abundance for *S. epidermidis* in most skin sites (Fig. 2C).

### Structural variants of *Staphylococcus epidermidis*

Community homeostasis is maintained by both host factors, interspecies interactions, and acquisition of genes, the latter of which can play important role in modulating adaptability to a particular environmental niche. These genes can encode for proteins that can influence signaling, virulence, and antimicrobial properties^16,17^. Consequently, we examined microbial structural variants that may potentially harbor genes that play a role in strain adaptability to a specific skin site. We focused our analysis on *S. epidermidis* because unlike other *Staphylococci*, it thrives well in multiple skin sites (Fig. 2A).

We retrieved pangenome sequences from panDB, a database which assembles non-redundant genomic regions from multiple sequenced strains^18^, split sequences into 1 Kb fragments, and determined the enrichment of fragments across samples. Because *Staphylococcus* other than *S. epidermidis* thrives mostly in foot sites, and less in sebaceous regions (Fig. 2A), we investigated structural variants that might be associated with *S. epidermidis* adaptability in sebum-rich sites. We identified 6 *S. epidermidis* fragments that were always enriched in sebaceous sites when compared with other sites (Fig. 3A) which suggests that these may harbor genes essential for survival in sebaceous regions.

**Figure 3.**
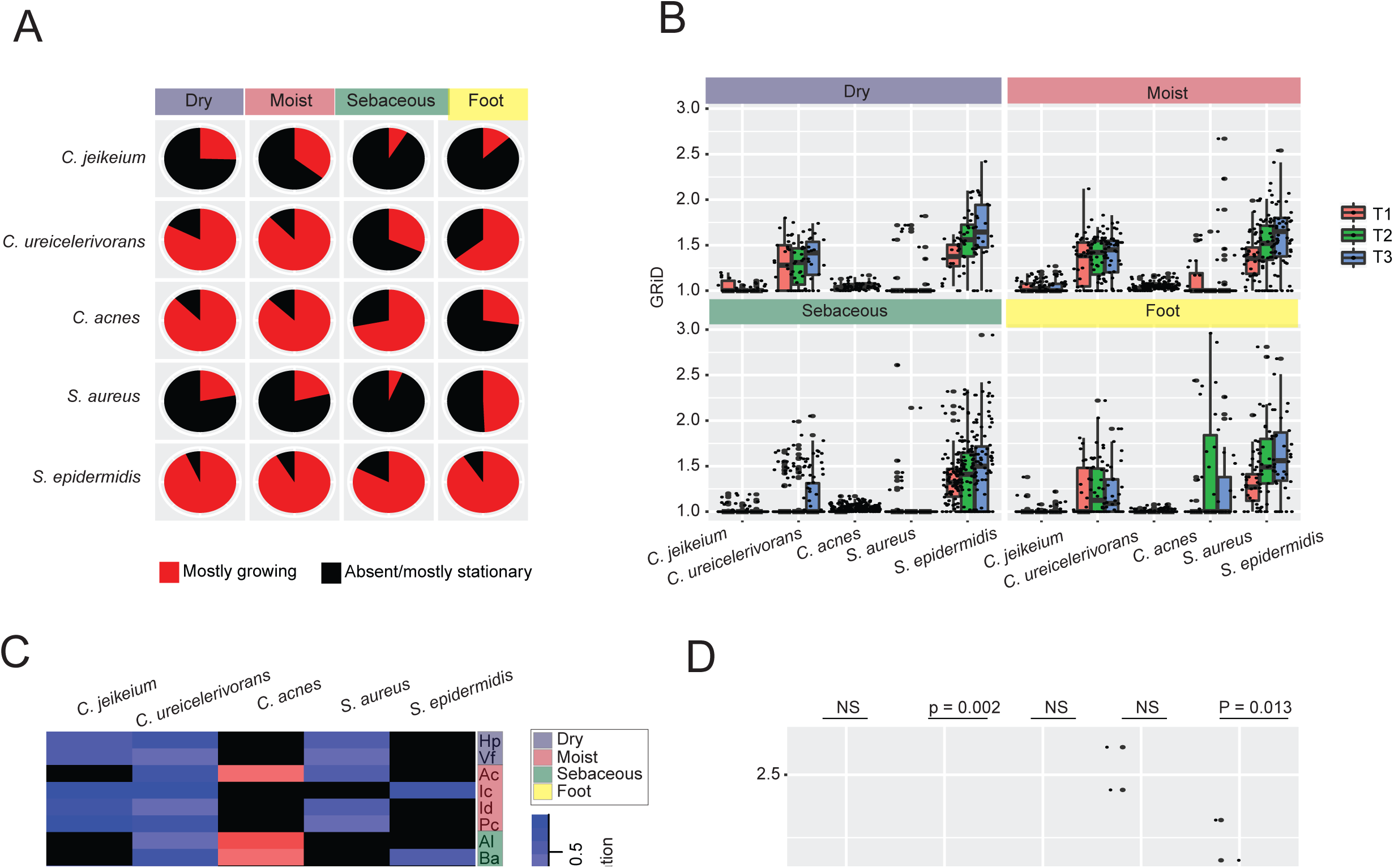
Structural variants in *Staphylococcus epidermidis*. **(A)** Venn diagram showing number of fragments differentially enriched in sebaceous sites when compared with dry, moist, or foot sites. **(B)** Phylogenetic tree constructed for genes identified in candidate fragments of *S. epidermidis* and *S. capitis.* Genes from both species clustering together are highlighted. **(C)** Candidate *S. epidermidis* fragment, corresponding gene, and functional annotation. **(D)** Spearman correlation between candidate *S. epidermidis* gene fragments and relative abundance or growth rate (GRiD) in sebaceous and foot sites.

In addition, we hypothesized that if these fragments are indeed associated with adaptability, homologues in *S. capitis*, a closely related genome, will also be associated with adaptability in the latter. Similar to *S. epidermidis*, we identified 11 *S. capitis* fragments that were always differentially enriched in sebaceous sites (Fig. 3A). Interesting, 4 of the 7 genes harbored in *S. epidermidis* fragments had homologues in *S. capitis* (Fig. 3B), suggesting an underlying importance in sebaceous sites.

Next, we examined if these candidate genes could be located in mobile genetic elements which may suggest inter/intra-species transmission. Using BLASTn alignment to the NCBI nt database, all candidate genes could be identified in previously annotated plasmids or phages (Fig. 3C). We conducted additional benchmark analysis to minimize false positives by determining the correlation between candidate genes and *S. epidermidis* abundance or growth rate. Unsurprisingly, all fragments containing genes with homologues in *S. capitis* were positively correlated with both abundance and growth rate in sebaceous sites. Most of these genes encode for hypothetical proteins; however, a candidate (S_epi_13619) encodes for a stress response protein suggesting that mechanisms to adapt to otherwise unfavorable conditions may play a important role in survival.

### Indirect interactions: horizontally-transferred genes in skin microbes

To further investigate the genetic basis of interspecies interactions using our refined metagenomic analysis, we investigated horizontal gene transfer (HGT). HGT is a mechanism by which a microbe can acquire genetic material that may confer an increased survival or competitive advantage within a community.

We developed a HGT prediction pipeline (Supplementary Fig. 5) and based on the simulated dataset, the pipeline was able to identify 31% - 51% of the simulated HGT genes. We found that the prediction sensitivity was confined by the sensitivity of the metagenomic assembler (i.e., whether a gene was fully assembled) and the sensitivity of the gene predictor (i.e., whether an open reading frame was correctly annotated as a gene) (Supplementary Fig. 6), but not the synonymous distance-based algorithm presented in this study. The predicted HGT genes exhibited a large variation of synonymous distances, representing both recent HGT events and more ancient HGT events during the diversification of the microbial species (Supplementary Fig. 6). Importantly, the HGT genes with the lowest synonymous distances (synonymous distance < 0.1), which correspond to the most recent HGT events, almost exclusively matched the simulated HGT genes (94% - 100%, Supplementary Fig. 6), demonstrating the high specificity of the prediction pipeline.

To predict HGT among microbes within the skin microbiome, we developed a pipeline using our existing metagenomic data. In each pair of assembled genomes, we identified HGT candidates by looking for pairs of genes that are significantly more similar than immobile genes (Fig. 4A, Supplementary Fig. 5). Our HGT identification pipeline is a parametric version of a previously described robust method that searched for identical or near-identical gene pairs in distantly related microbial genomes^19^. Consistent with previous reports, functional annotation of HGT candidates showed a distribution over a wide functional spectrum^19^ (Fig. 4B). The most overrepresented functionality of the predicted HGT candidates were the transporters, highlighting the potential of the microbes to acquire the ability to uptake environmental nutrients and extrude harmful molecules through HGT events (Fig. 4B). Although most types of transporters were uniformly distributed in the mobile gene pool, microbiome at the sebaceous sites demonstrate enrichment of transporters that are involved in transporting metallic cation (Fig. 4B and 4C), including iron, manganese, zinc, cobalt, nickel, and biotin. These results suggest the existence of a pool of (conditionally) mobile genes exerting a multitude of biological functions, with enrichment of specific functions observed at specific skin types.

**Figure 4.**
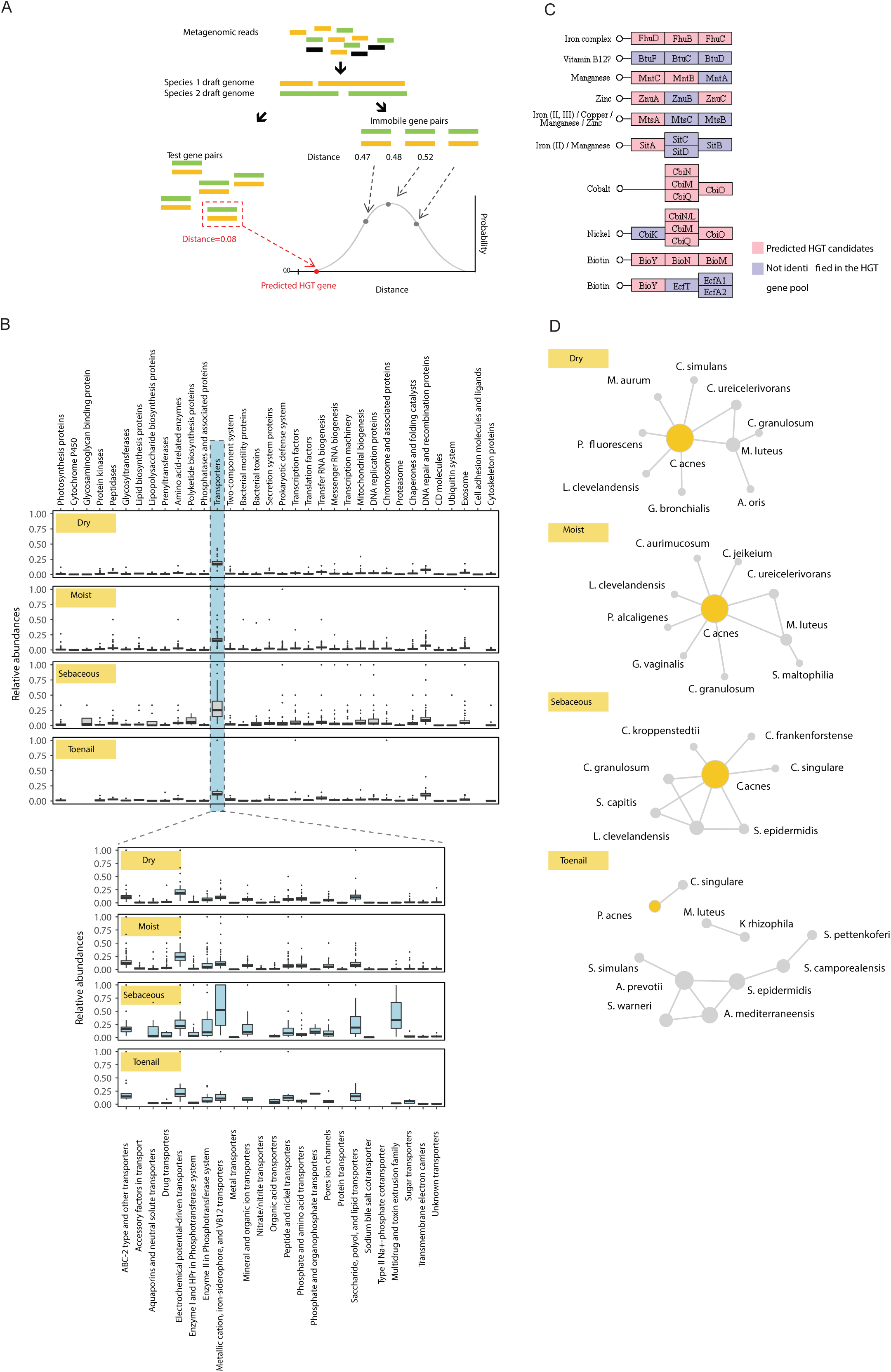
Horizontal transfer of genes in skin community. **(A)** Overview of the HGT candidate identification process. Metagenomic reads were assembled and contigs belonging to the same species were pooled into a species draft genome. For each pair of species draft genomes, orthologous gene pairs were predicted. If a gene pair had a significantly smaller synonymous distance than the immobile gene pairs (that is, the universal single-copy orthologs), the pair of genes were identified as HGT candidates. **(B)** Distribution of functions (i.e., KEGG BRITES annotations) of all identified horizontal gene transfer (HGT) candidates and candidates that were annotated as transporters. **(C)** Detailed functions (i.e., KEGG orthologs) of HGT candidates annotated as metallic cation, iron-siderophore, and vitamin B12 transporters identified in the sebaceous sites. **(D)** Networks representing the top 10 species pairs for which HGT events were most frequent (i.e., HGT events were identified in the largest amount of samples) in each type of skin site. Nodes represent species and edges represent HGT events. In each type of skin site, the size of a node is proportional to the degree of that node.

Finally, we constructed a network to reflect the top HGT events among microbial species in each type of skin sites (Figure 4D). Across skin sites, HGT were identified as a function of abundant species. For example, HGT candidates identified at a sebaceous site predominantly came from *Actinobacteria*, including bacteria in the genera *Propionibacterium, Corynebacterium* and *Staphylococcus* (Fig. 4D). Due to its dominance in many skin sites, *C. acnes* was central in networks corresponding to all types of skin types except for toenails, in which *Actinobacteria* (*C. acnes, Corynebacterium singulare, Micrococcus luteus*, and *Kocuria rhizophila*) and multiple *Staphylococcus* species formed disconnected networks, strongly suggesting that HGT events at a body site were driven by the microbiome composition at the site. Overall, this pipeline characterizes statistically likely HGT candidates directly from shotgun metagenomic data, and is useful to simultaneously estimate the functional and taxonomic distribution of the mobile genes between genera.

## DISCUSSION

The human skin, our largest organ and first line of defense, is home to a diversity of microorganisms. These microbes play an essential role in influencing metabolic processes, immune system modulation, and antagonism of potentially transient pathogens^15^. Alterations in community composition have been associated with a number of skin diseases like atopic dermatitis, psoriasis, and eczema^20,21^. Numerous host-intrinsic (e.g., genetics, immune competence, skin barrier conditions), environmental or lifestyle factors (e.g., hygiene, exposure to different microbes), as well as microbiome-intrinsic factors affect disease predilection. A deep understanding of these factors, as well as the ecological complexity of the skin’s microbiota, is needed to understand factors that influence its homeostasis and predisposition to disease.

Large-scale studies have aimed to characterize the skin microbiota using deep shotgun metagenomic sequencing, yet conclusions drawn from those previous analyses were limited in that a majority of sequence reads could not be mapped to any known genome^5,6^. To solve this limitation, we used an integrated approach that incorporated *de novo* assembly and binning with reference-based analyses to identify and quantify new microbial skin occupants. This dual approach provided a key improvement in the resolution of the community. While we observed a significant reduction in uncharacterized sequence space using *de novo* approaches, viral genomes and low abundance genomes were poorly captured but could be resolved with reference genomes.

High-level analyses of the defining characteristics of the skin microbiome were largely concordant with previous findings. However, our hybrid approach provided important new insights into skin community structure, including: increased diversity, reduced stability across many sites, dominance of previously uncharacterized microbes in certain skin sites like the external auditory canal, and reduced representation of phage than previously believed. With few exceptions, we found that previous reports overestimated stability of the skin microbiome, likely because they lacked deeper classification of additional genomes, particularly for staphylococci and cornyeforms. Subsequently, we focused our analysis—continuing to interleave reference-based and *de novo* approaches—to examine potential factors that could underlie community stability and homeostasis, including the growth rate of different community members and interspecies interactions.

Leveraging metagenomic data to predict growth rate of dominant skin species, we measured marked variance in which species were actively growing in the skin, and how skin site could affect activity. For example, most microbes in dry sites were actively dividing compared to relatively few in sebaceous sites. Moreover, *S. epidermidis* appeared to grow exponentially in all sites, at similar growth rates, suggesting that the skin environment generally provides adequate nutrients to support its rapid growth. Alternatively, *S. epidermidis* strains may have acquired specific genes modulating adaptability to each site. For example, we identified numerous genes present in mobile genetic elements that were associated with adaptability to sebaceous sites. Yet, strikingly, its ultimate relative abundance was not correlated with growth rate in most skin sites, suggesting an equally rapid killing or cell death. In this case, the lack of correlation between growth rate and relative abundance indicated that competitiveness within the community is largely independent of growth rate. Other skin sites/species showed a different relationship between growth rate and relative abundance. A positive correlation suggests that strains out-compete the rest of the community during exponential growth, as in the case of *C. ureicelerivorans*. Conversely, a negative correlation may reflect an active quorum sensing mechanism involved in the regulation of growth rate. For example, we observed a negative correlation for *C. acnes* that is restricted to sebaceous skin, suggesting that the microenvironment may play a role in regulating growth rate.

In addition, we observed that several types of transporters, especially the metal transporters, were highly abundant in the HGT gene pools at the sebaceous sites, potentially reflecting the importance to transport a vast variety of lipid soluble metals diffusing through the permeable cells of the sebaceous glands and follicular walls – a major absorption pathway of metals, including metallic toxicants. Additionally, the enrichment of metal transporters at the sebaceous sites paralleled the over-representation of *Actinobacteria* species at those skin sites, raising the possibility that the dissemination of metal transporting ability among Actinobacteria microbes at the sebaceous sites may be important to metal balance and consequently the health of the host.

In conclusion, we present a new landscape of the skin microbiome, providing the highest resolution reconstruction of microbial community composition and biodiversity in the skin to date. Importantly, we have used new approaches and analyses in a multifaceted investigation of functional elements and interspecies interactions underlying stability and community homeostasis. Our findings have generated testable hypotheses to interrogate interspecies and/or interstrain inhibition. Finally, our analyses are broadly applicable to investigate these mechanisms in skin disease, which ultimately can provide clues as to sources of antimicrobials directed towards strain-specific pathogens.

## METHODS

### Sample datasets

We retrieved 698 metagenomic shotgun skin samples from our previous work^5,6^, which have been quality filtered for the presence of human DNA. The majority of the samples (n = 594) in these dataset were derived from longitudinal sampling of 12 individuals at 3 different time points with sampling intervals of 10-30 months (“long”) and 5-10 weeks (“short”). 24 samples from this collection were also derived from 2 individuals with hyper IgE syndrome. The remaining samples represent a single timepoint from three additional healthy individuals.

### Taxonomic classification of skin microbes

To classify skin microbes, we used a hybrid *de novo*-based and referenced-based characterization technique (Supplementary Fig. 2). All samples were concatenated and sequence reads were assembled into contigs using MEGAHIT 1.0.5 (--k-min 37 --k-max 67 --k-step 10 -m 0.99 --kmin-1pass --continue)^11^, which was used for its ability to handle large datasets with relatively low memory requirement and short run time. We discarded contigs < 1Kb, mapped each individual sample’s reads back to the contig catalog using bowtie2 2.2.8 (--sensitive)^22^, and extracted unmapped reads. However, read coverage from our derived contigs catalogue was relatively low (Supplementary Fig. 2). To re-assemble unmapped reads, we concatenated a subset of unmapped reads from randomly selected samples (due to the high memory requirement of SPAdes^23^) and re-performed previous steps using SPAdes 3.7.1 (--meta) for *de novo* assembly, which was better able to resolve scaffolds from contigs. Newly extracted contigs/scaffolds were merged with the previous catalog to produce a non-redundant contigs catalog. We repeated the mapping of reads to the catalog to retrieve unmapped reads, randomly concatenated a small subset of unmapped reads, and performed assembly using SPAdes. We repeated this iterative step until no additional improvement in reads coverage from our contigs/scaffolds catalog was observed (Supplementary Fig. 2), obtaining a total of 1,037,465 contigs/scaffolds > 1Kb.

Contigs/scaffolds were then grouped into genome bins using MetaBAT (--sensitive, -m 1500)^24^, which resulted in 556 bins. Bins were taxonomically classified using MEGAN 4.70.4^25^ (Supplementary Table 1). We excluded 22 bins which were of non-microbial origin and further evaluated the quality of each bin using CheckM 1.0.6^26^. For stringent annotation, we required that ≥ 65% of contigs/scaffolds present in a bin were assigned to the lowest level taxonomy; the sole exception being the kingdom-level taxonomy where our requirement was 40%. Bins were labeled “uncharacterized” if they were unable to be assigned to at least a kingdom, although by further relaxing our requirements to exclude contigs/scaffolds with no hits enabled characterization of those bins to at least the kingdom level (Supplementary Fig. 3B).

Each sample was then mapped back to the genome bins using bowtie2. Unmapped reads were subsequently characterized by a reference-based approach using reprDB^18^ and assigned to a genome using Pathoscope 2.0^8^. Reads unassignable by either method were categorized as “dark matter”.

### Bacterial growth rate estimation from skin samples

We estimated bacterial growth rate using GRiD (v1.3)^13^ using a coverage cutoff of 0.2. We created custom GRiD database using species of *C. acnes, S. epidermidis, S. aureus, C. jeikeium*, and *C. ureicelerivorans*.

### Structural variant analysis

We retrieved pangenome sequences for *S. epidermidis* and *S. capitis* from panDB^18^, split genomes into 1 Kb fragments, and predicted genes using prokka^27^. We mapped reads using bowtie2 to the fragment pool of each species, estimated reads count across samples using featureCounts^28^, and DESeq2^29^ to infer genes differentially enriched between skin sites. We filtered candidates using a adjusted p-value of 0.05 and log_2_ fold change of 1. We aligned genes from *S. epidermidis* and *S. capitis* to generate a phylogenetic tree using MAFFT^30^.

### Horizontal gene transfer prediction

By definition, HGT genes that transferred between two lineages have shorter evolutionary distance than immobile genes (those that diverged at the time of divergence of the lineages). Therefore, HGT genes tend to be more similar^19^. For a pair of genomes, one reliable method of identifying HGT genes is to search for pairs of genes that are more similar to each other than the immobile genes in the genomes that reflect the evolutionary distance of the genomes. Genomes assembled from metagenomic shotgun data are inevitably incomplete and inaccurate, especially at strain level. Therefore, we assembled genomes at species level and proceeded only with genomes that have at least 25% completeness. To do this, metagenomic shotgun reads were first assembled using MEGAHIT. The resulting contigs were then assigned a taxonomic label using Kraken v0.10.6^31^; all contigs assigned to the same species were combined to represent the species draft genome. Genes were predicted from each species draft genome using prodigal^32^, and completeness of the assemblies assessed using BUSCO v2^33^.

We identified potential HGT genes in each pair of species genomes. To do this, we first assembled a set of immobile genes from the genome pair to compute a null distribution of sequence similarity. Immobile genes were identified using the bacteria-specific universal single-copy orthologs (USCOs) annotated by BUSCO. Because USCOs are universally present across bacterial lineages and exist only in single copy, their horizontal transfer is unlikely. Second, all gene sequences from a pair of species genomes were clustered using uclust^34^ at 0.5 similarity cut-off to reduce complexity. If any pair of genes within a cluster and from different species genomes has a significantly higher similarity than the immobile genes, the gene pair is identified as horizontally transferred. To mitigate the effect of natural selection, we computed synonymous distance—the number of synonymous changes per synonymous site—as the test statistics for similarity. Each pair of protein-coding genes was first reverse-aligned using the seqinr package^35^, after which synonymous distance was computed using PAML^36^, which implements an *ad hoc* method that corrects for codon frequency bias. Finally, to further lessen the influence of purifying selection, we removed HGT candidates that represented essential genes (i.e., genes with positive hits to the DEG 10 database^37^ using ublast^34^ with e-value < 10^−9^) and ribosome genes (ie, genes corresponding to KEGG BRITE ko03009 and ko03011).

We applied the prediction pipeline to three simulated microbial communities for validation. The simulated communities was generated using HgtSIM^38^. Briefly, each community included three common skin bacteria species: *C. acnes, S. epidermidis*, and *Streptococcus mitis*. Each species was represented by three sequenced strain genomes, including one RefSeq representative strain and two other strains from RefSeq. All 9 strains were mixed at equal abundances in each simulated community. From the representative strain, 5 genes were randomly selected and horizontally transferred to all other strains in the community (a total of 15 HGT genes in each community). The three microbial communities differ in the amount of mutations accumulated in the HGT genes: the HGT genes were allowed to accumulate 0, 5% and 10% mutated bases in the recipient strains for the three communities, respectively. 3 million paired-end metagenomic shotgun reads were sampled from each community as part of the HgtSIM pipeline, and subsequently processed using the HGT prediction pipeline described above.

### Statistics

We conducted all statistical analyses using R software. Spearman correlation was utilized for all correlation coefficient analyses while statistical differences between population groups were determined using the Wilcoxon rank-sum test. Where multiple measurements (e.g., timepoint, skin site within an individual) were used for correlation analyses, partial Spearman correlation was used, adjusting for multiple measurements. To assess microbial stability over time, we utilized the Yue-Clayton theta index, which calculates the distance between communities based on relative abundance of species in the population^39^. Community diversity was determined using the Shannon diversity index, which measures both species richness and evenness.

## DECLARATIONS

### Ethics approval and consent to participate

Not applicable.

### Consent for publication

All authors have approved submission of this manuscript.

### Availability of data and material

The data used in this analysis is available in SRA under Bioproject 46333.

### Authors’ contributions

AE and JO conceived the project. WZ contributed scripts and analysis. AE and JO analyzed data. AE and JO wrote the manuscript.

## Acknowledgements

We would like to thank the Oh lab for critical reading of the manuscript.

## SUPPLEMENTARY INFORMATION

**Supplementary Figure 1.**
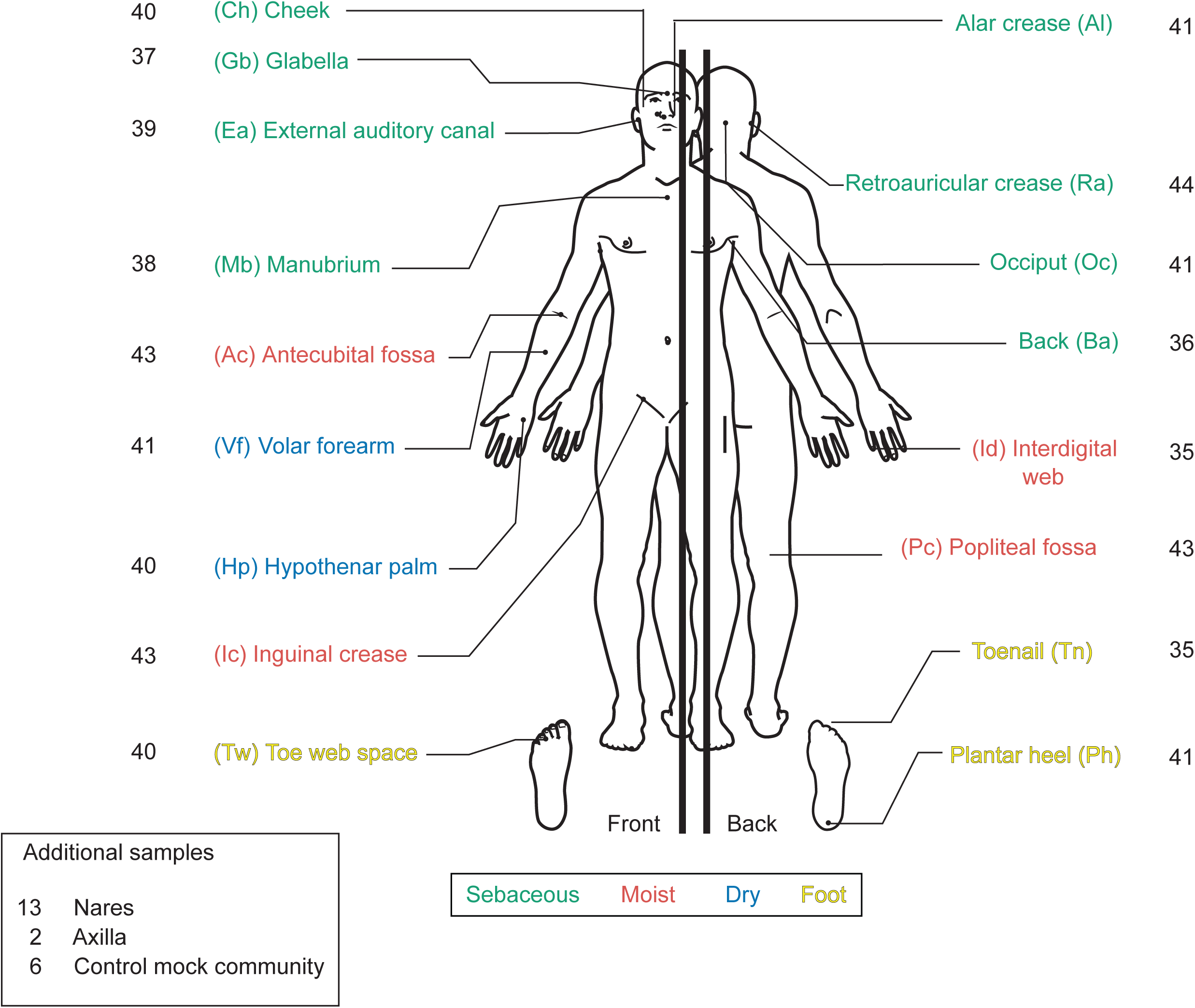
Skin sites, skin physiologic characteristics, and number of samples used. Overview of 698 samples analyzed in this study, encompassing 15 healthy adults and two hyper-IgE patients, three timepoints, and 17 skin sites, representing 4 microenvironments; dry, moist, sebaceous, and foot sites. The numbers adjacent each site correspond to the total number of samples derived from those sites.

**Supplementary Figure 2.**
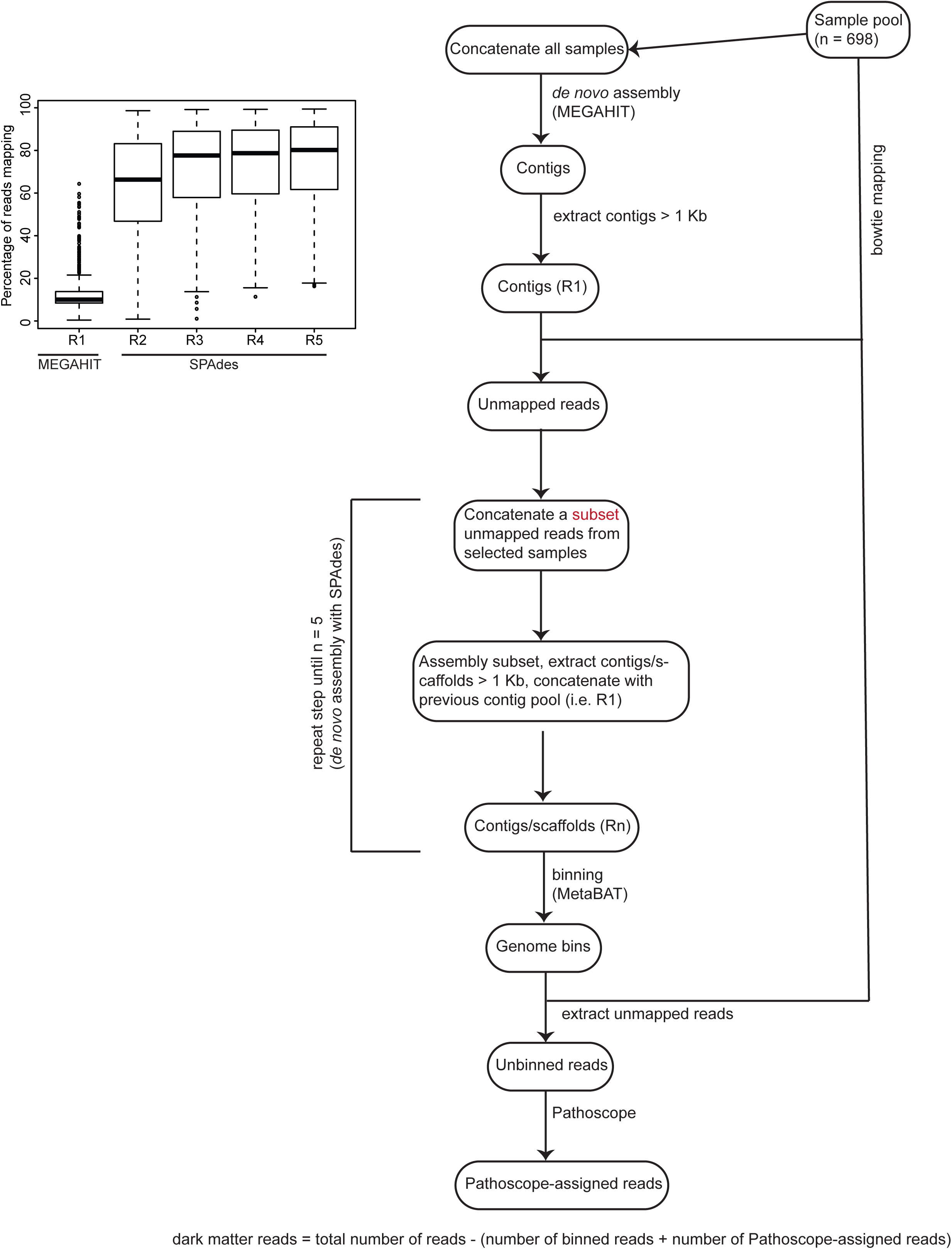
Flowchart of hybrid *de novo* and reference-based approach to resolve microbial dark matter. The inset boxplot represents the percentage of reads mapping back to contigs/scaffolds (> 1 Kb) catalog after each round of iterative assembly. Black lines indicate median; boxes first and third quartiles.

**Supplementary Figure 3.**
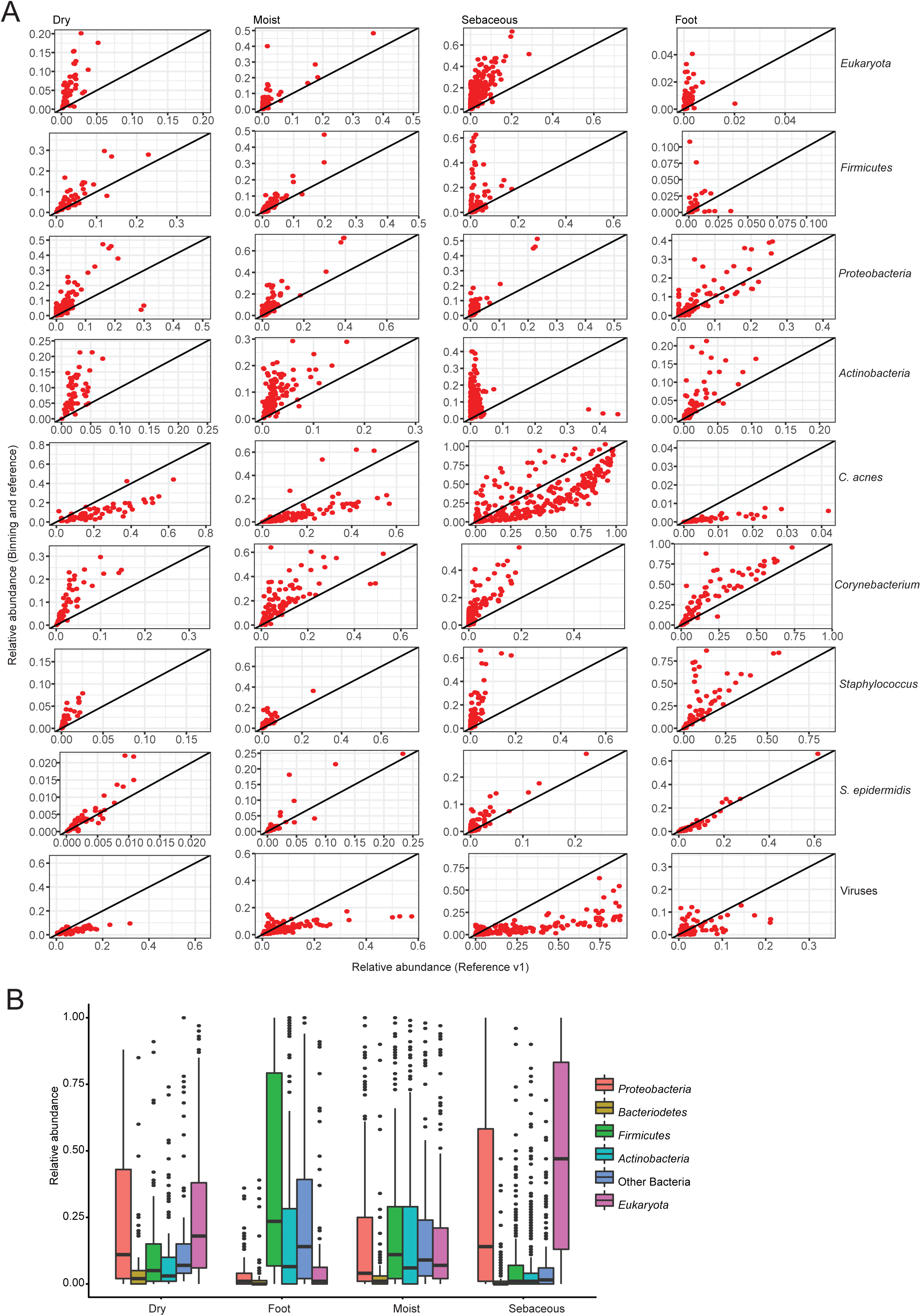
Hybrid *de novo* and reference-based resolution of dark matter. **(A)** Scatterplots show concordance in relative abundance between the original classification in Oh *et al*.^6^ and the hybrid approach. Points deviating significantly from the diagonal are those whose relative abundance changed significantly based on the resolution of dark matter. Major genera, phyla, and kingdoms are shown. **(B)** Annotation of uncharacterized bins. Boxplots shows proportion of uncharacterized bins reassigned to a taxa. We relaxed our initial annotation requirement by excluding contigs/scaffolds with no hits from MEGAN output and re-ran our annotation pipeline.

**Supplementary Figure 4.**
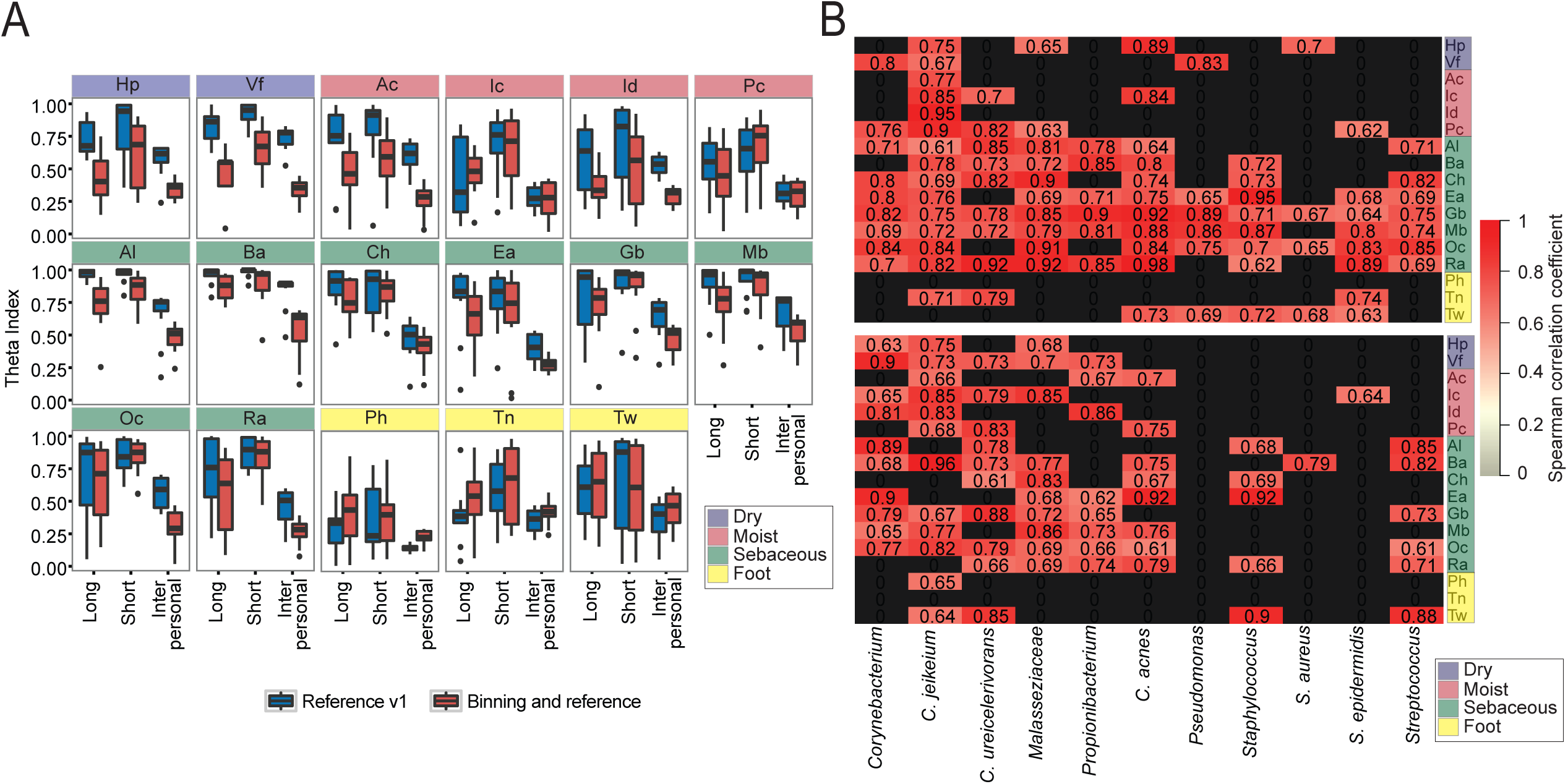
Higher compositional reconstruction of the skin microbiome. **(A)** Community stability across skin site as calculated by the Yue-Clayton theta index, where θ∼1 represents a completely stable community. “Long” refers to sampling time interval between T1 and T2 while “Short” represents short sampling time interval between T2 to T3. **(B)** Heatmap shows partial Spearman correlation values correcting for multiple measurements between microbial relative abundance at timepoints T2 vs T3 (i.e. short time interval) (top) and T1 vs T2 (i.e. long time interval) (bottom). Black colors indicate no correlation. *C. acnes=Cutibacterium* acnes, C.=*Corynebacterium, S.*=*Staphylococcus*.

**Supplementary Figure 5.**
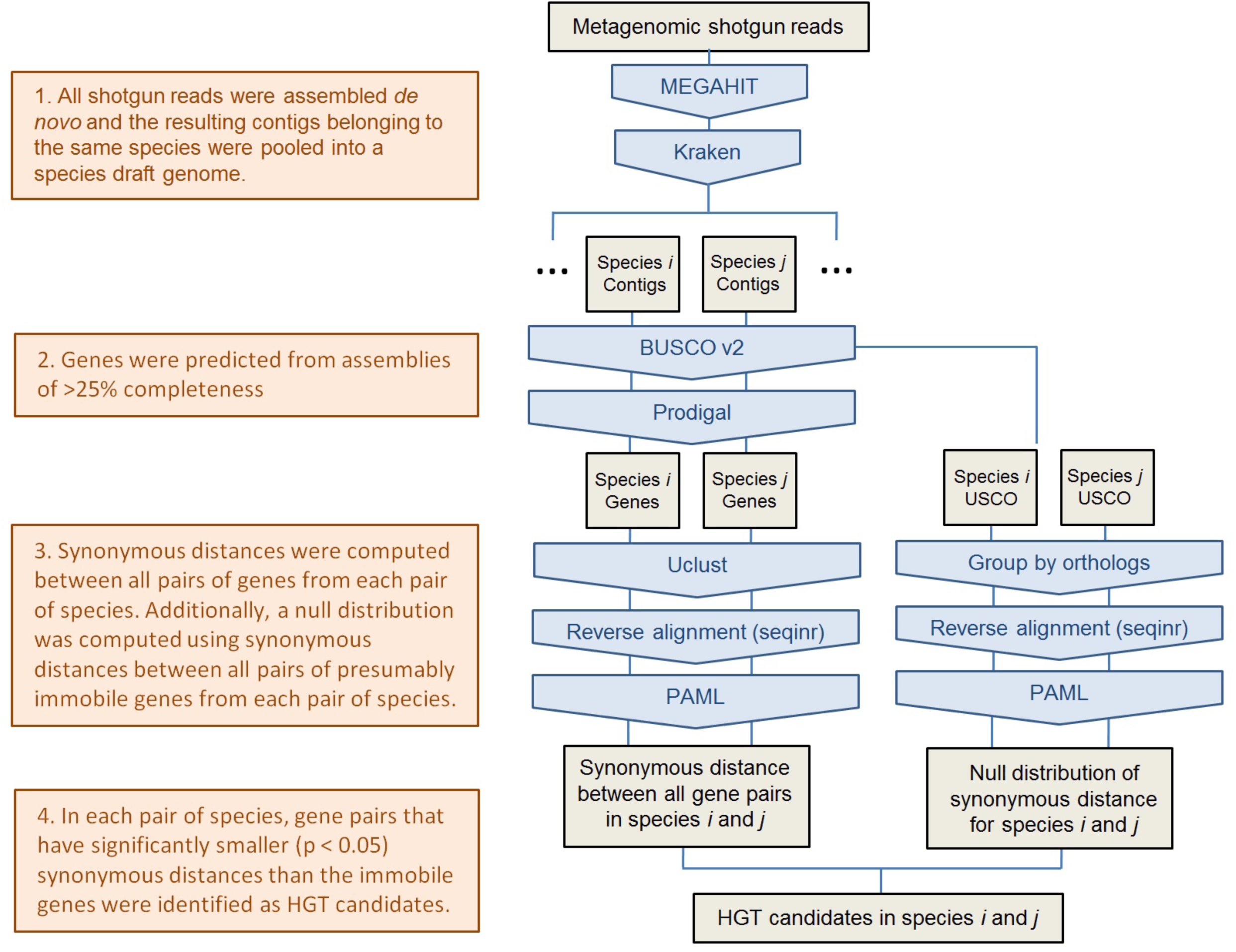
HGT candidate identification pipeline.

**Supplementary Figure 6.**
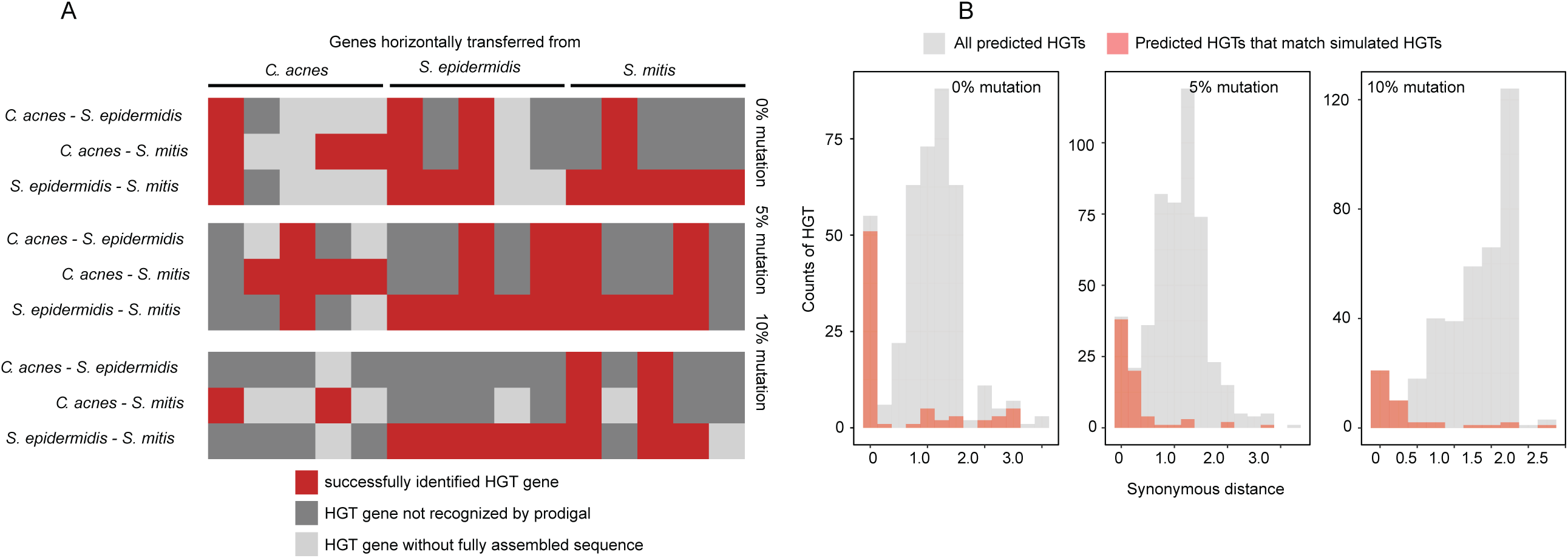
Validation of the HGT prediction pipeline. **(A)** Simulated HGT genes that were identified using the pipeline between all species pairs in all simulated communities (0%, 5%, and 10% mutations). **(B)** Number of HGT genes identified as a function of the synonymous distance of the HGT genes in the species pairs. Identified HGT genes that matched the simulated HGT genes were shown in red.

**Supplementary Table 1:** Microbial genome bins identified using *de novo* approach. Bins highlighted in yellow represent non-microbial genomes.

**Supplementary Table 2:** Bin coverage across samples.

**Supplementary Table 3:** Community relative abundance resolved using hybrid *de novo* and reference-based approach

